# Quantifying Cellular Pluripotency and Pathway Robustness through Forman-Ricci Curvature

**DOI:** 10.1101/2021.10.03.462918

**Authors:** Kevin A. Murgas, Emil Saucan, Romeil Sandhu

## Abstract

In stem cell biology, cellular pluripotency describes the capacity of a given cell to differentiate into multiple cell types. From a statistical physics perspective, entropy provides a statistical measure of randomness and has been demonstrated as a way to quantitate pluripotency when considering biological gene networks. Furthermore, recent theoretical work has established a relationship between Ricci curvature (a geometric measure of “flatness”) and entropy (also related to robustness), which one can exploit to link the geometric quantity of curvature to the statistical quantity of entropy. Therefore, this study seeks to explore Ricci curvature in biological gene networks as a descriptor of pluripotency and robustness among gene pathways. Here, we investigate Forman-Ricci curvature, a combinatorial discretization of Ricci curvature, along with network entropy, to explore the relationship of the two quantities as they occur in gene networks. First, we demonstrate our approach on an experiment of stem cell gene expression data. As expected, we find Ricci curvature directly correlates with network entropy, suggesting Ricci curvature could serve as an indicator for cellular pluripotency much like entropy. Second, we measure Forman-Ricci curvature in a dataset of cancer and non-cancer cells from melanoma patients. We again find Ricci curvature is increased in the cancer state, reflecting increased pluripotency or “stemness”. Further, we locally examine curvature on the gene level to identify several genes and gene pathways with known relevance to melanoma. In turn, we conclude Forman-Ricci curvature provides valuable biological information related to pluripotency and pathway functionality. In particular, the advantages of this geometric approach are promising for extension to higher-order topological structures in order to represent more complex features of biological systems.

## 1 INTRODUCTION

A major biological phenomenon is cellular differentiation, whereby pluripotent stem cells undergo phenotypic evolution to approach one of many possible differentiated states. Understanding how cellular dynamics shape the trajectories of differentiation can provide us with a deeper knowledge of the processes that govern phenotypes, which could allow us to predict cellular behavior in various biological processes and ultimately may inform our strategy for influencing these complex biological systems with targeted pharmacology. For example, recent work in this regard have ranged from stem cell biology [1–4], cancer biology [1, 2], cellular reprogramming [5], to defining biological robustness and drug resistance [6–8]. Epigenetic evolution is familiarly motivated by C.H. Waddington [9] as a hill-like “landscape”, in which cellular state is intrinsically coupled to a quantity similar to potential energy in classical mechanics (Fig. 1A). That is, cellular differentiation could be modeled as a ball rolling down the hillside of this landscape, gaining momentum and following “grooves” towards a state of lower potential energy and perhaps settling into a local minima. However, the mathematical quantification of this landscape in regards to cellular trajectories and pluripotency in general has only been recently investigated [3, 4, 10] and to a large part, remains an open problem.

**Fig. 1.**
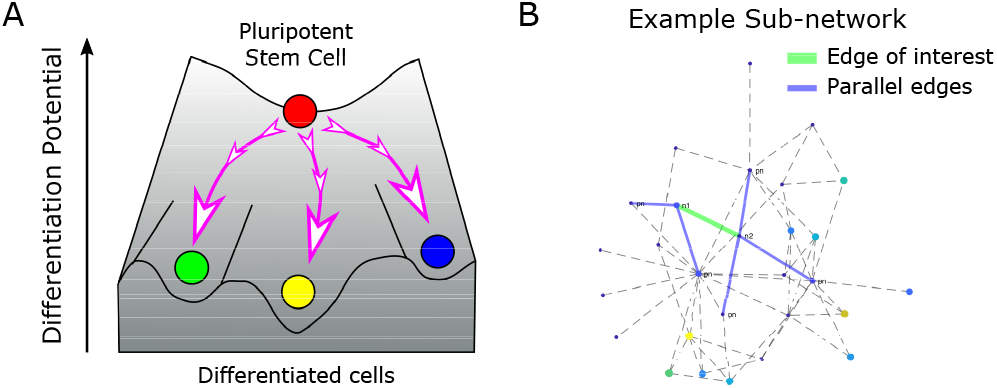
A: Stem cell differentiation as a landscape, wherein stem cells move down a gradient of potential energy to reach local minima at unique differentiated cell states. B: Example subnetwork of PPI network, with an edge of interest (green) and corresponding parallel edges (blue) labelled.

To this end, in order to estimate and capture the complex dynamics of cellular differentiation, one can model the underlying biological systems as a network. While a complete review of such network modeling is beyond the scope of this work [11], this study focuses on protein-protein interaction (PPI) networks [12, 13]. Here, individual genes are treated as components in a large network of interacting proteins, wherein relationships such as protein binding or enzyme catalysis are modeled as corresponding edges between nodes on a graph and for which such “interactions” have been validated by experimental studies. In this regard, many recent studies have applied PPI networks to model the differentiation landscape [1, 2, 14]. More importantly, several of these recent works attempt to mathematically quantify the general notion of pluripotency from a statistical mechanics perspective, in which the differentiation capacity of a cell can be described by entropy measured on the PPI network. In particular, [14] utilizes a notion of graph entropy to examine the “randomness” of gene expression data overlaid onto a PPI network. These studies demonstrate gene network entropy to be a powerful statistical descriptor of cell pluripotency, decreasing upon cellular differentiation and increasing in situations such as cancer [1, 2].

While statistical mechanics and in particular notions of entropy are well-known with ubiquitous applications beyond cellular biology, recent work has shown that Ricci curvature (a geometric measure of “flatness”) may also be reinterpreted as a statistical quantity and proxy for entropy [15–17]. This is particularly fascinating in the continuous case as one is able to unify a purely deterministic geometric (local) quantity of curvature with that of a statistical (equilibrium) quantity of entropy. From the biological perspective, this provides a “bridge” to investigate geometric (and topological) properties of biological networks to better capture cellular functionality in facets not generally realizable from a purely classical statistical lens [18]. That said, discretizing the continuous definition of Ricci curvature (and other forms of curvature in Riemannian geometry [19]) is not necessarily straightforward especially over a discrete metric space (e.g., graph) where differential operators are not readily provided. As such, several works have focused on discretizing Ricci curvature for graphs including the definitions of Ollivier [20] and Forman [22]. While both discretize Ricci curvature for graphs, Ollivier-Ricci curvature [20] is motivated through the statistical idea that probabilistic neighborhoods are closer (farther) than their centers depending on positive (negative) curvature, using optimal transport theory to compute such a distance, whereas Forman-Ricci curvature provides an explicit combinatorial curvature formula based on Bochner’s method of decomposing the combinatorial Laplacian (more detail in Section 2) [22–25]. While both of these measures work well on 1-dimensional simplicial complexes, i.e. graphs with only nodes and undirected edges, Forman-Ricci curvature is in fact formulated for higher order structures such as faces, which are composed of multiple nodes and edges [26]. Therefore, in the context of biological gene networks where multiple genes can interact simultaneously in common pathways, Forman-Ricci curvature may prove advantageous over previous applications of Ollivier-Ricci curvature from both computational and mathematical perspectives.

Of importance, theoretical work [15–17] has demonstrated a relationship between Boltzmann entropy and Ricci curvature (i.e. changes in entropy are positively correlated with changes in curvature). Several works have exploited this concept in a variety of venues from cancer biology [18, 27, 28], neuroscience [29, 30], wireless network congestion [31], economics [32], to graph-based Ricci flows [25, 33]. Here, most of the biological work has focused on Ollivier’s definition. However, there exist intrinsic computational and mathematical limitations to the generalization of this definition to the higher order topological structures necessary to capture biological functionality underlying cellular dynamics. Therefore, our study examines curvature and entropy in the context of stem cell differentiation and cancer with the goal of establishing a direct link between the deterministic quantity of Forman-Ricci curvature and the statistical quantity of network entropy. Ultimately, we aim to explore the capability of Forman-Ricci curvature as a meaningful descriptor of cellular pluripotency and pathway functionality. If curvature analysis can indicate which genes are associated with pluripotency and how these genes drive pluripotency, this would suggest geometric theory could guide applications in biology and medicine to exert dynamic control over cellular differentiation processes and pathologic processes as in cancer. Perhaps more importantly, this will lay the foundation for an exploration of higher-dimensional biological structures through extension of Forman’s definition.

This paper is organized as follows: In Section 2, we provide preliminaries and necessary background. In Section 3, we describe our computational approach. Following this, Section 4 presents results that highlight Forman-Ricci curvature as a descriptor of pluripotency in a stem cell dataset and pathway functionality in a melanoma dataset. Lastly, Section 5 presents conclusions and discussions of future directions.

## 2 BACKGROUND

### 2.1 Motivation: Interplay of Entropy & Curvature

Critical to our motivation for this study are several works exploring the relationship of entropy and curvature in geometric spaces [15–17]. The core of the approach of these studies involves the dependence of various geometric quantities on the lower bound of Ricci curvature. Namely, for a given Riemannian manifold, the existence of a lower quadratic bound on Ricci curvature, *Ric*(*M*) ≥ *κI* where *I* is the identity matrix, implies K-convexity of Boltzmann entropy on the same manifold [17]. This property can be exploited to establish a positive correlation between changes in entropy Δ*S* and changes in curvature Δ*Ric*,

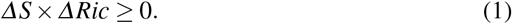

An additional key relationship to these two variables involves robustness, which can be defined as the rate of return to equilibrium in a dynamical system. The Fluctuation theorem [27, 34] asserts a positive correlation between changes in entropy Δ*S* and changes in robustness Δ*R*, which therefore, by Eq. (1), implies the same relationship between Ricci curvature and robustness,

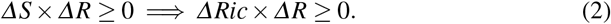

These findings provide direction for this study, serving as the theoretical groundwork for exploring Ricci curvature in gene networks as a statistical indicator for pluripotency and pathway robustness. Given that global network entropy has been demonstrated as a descriptor of cellular pluripotency [1, 2, 14], we aim to extend Ricci curvature as a geometric quantity on biological gene networks. Then, we can apply the relationships of curvature to entropy and robustness, respectively, to assess Ricci curvature as a potential descriptor of pluripotency (globally) and pathway robustness (locally).

### 2.2 Biological Graphs: Topological & Geometric Information

In the context of biology, graphs such as protein-protein interaction networks can represent the complex system of interactions between diverse proteins. On one hand, topological information alone can provide valuable information about biological networks [35]. Nevertheless, gene expression array data can be overlaid onto these graphs to augment the underlying topology with geometric node and edge weights, providing richer and more complex analysis than merely topological information. In this paper, we seek to analyze such gene networks from a geometric perspective.

Previous studies examining gene network entropy utilize a formulation based on Shannon entropy [14]. In brief, local entropy *S_i_* at a node *i* can be determined based on the interaction probabilities *p_ij_* of neighboring nodes, and a subsequent global entropy rate *SR* can be computed as a weighted-average of local entropies based on the stationary (invariant) distribution *π* of the network,

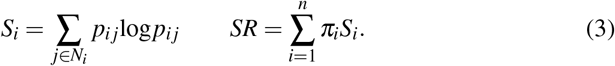

By realizing a gene network as a geometric object, we aim to exploit the relationship of entropy and Ricci curvature, shown in Eq. (1), to pivot from the statistical quantity of entropy into the geometric quantity of curvature. We also seek to utilize the relationship of robustness and Ricci curvature, in Eq. (2), by examining Ricci curvature within local network components, which here would represent individual genes and gene pathways, to quantify changes in functional robustness that could indicate critical pathways involved in biological processes such as tumorigenesis.

Importantly, the discrete structure of gene networks requires a discretization of Ricci curvature, for which we select Forman-Ricci curvature due to its combinatorial nature, computational advantages, and adaptability to higher-order structures.

### 2.3 Forman-Ricci Curvature

In the discrete setting, Forman-Ricci curvature can be developed through a combinatorial analogue to the Bochner-Weitzenböck decomposition of the Riemannian (or Hodge [21]) Laplacian □_*p*_ into a rough (or Bochner) Laplacian *B_p_* and a combinatorial curvature operator *F_p_* [22],

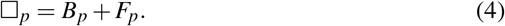

This definition applies to CW-complexes, a general class of topological structures that includes graphs. In this sense, a graph is considered a complex consisting of nodes (0-cells) and edges (1-cells) glued together at their boundaries, i.e. their nodes. For each edge in the graph, we use Forman’s approach to derive an explicit combinatorial curvature formula that depends only on the weights of the edge, the edge end-nodes, and neighboring (parallel) edges [22].

Let us consider an arbitrary graph consisting of nodes and edges (Fig. 1B). For a given edge *e* with ascribed edge weight *w_e_*, coupled with end-nodes *v*_1_ and *v*_2_ with respective node weights *w*_*v*_1__ and *w*_*v*_2__, Forman-Ricci curvature is defined as

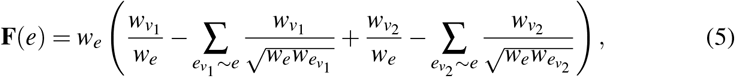

where *e_v_i__ ~ e* denotes parallel edges that share a common node *v_i_* with edge *e*.

This definition applies to both undirected and directed graphs [24]. In the case of a *directed* graph, the same definition is applied, with the caveat that only parallel edges concordant in direction with edge *e* are considered and any opposite edge 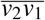 is disregarded. Notably, as we consider here only simple graphs of nodes (0-cells) and edges (1-cells), we disregard higher order structures such as faces (2-cells), which do appear in the full derivation [22].

As Forman-Ricci curvature is defined on each edge of the graph, one can define a contracted nodal curvature as the mean of all edge curvatures incident to a given node,

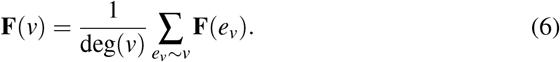

In the case of a *directed* graph, nodal curvature is defined as the mean of all incoming edge curvatures minus all outgoing curvatures,

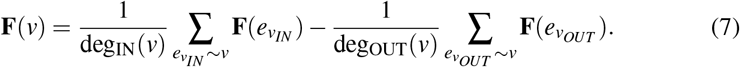

A global average of curvature **F***_GA_* can be computed as a weighted average of nodal curvatures, weighted by the stationary distribution *π* of the graph,

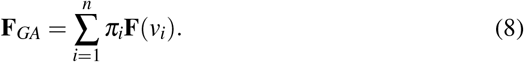

## 3 APPROACH

### 3.1 Gene Expression Pre-Processing to Graph Construction

In this note, we regard an RNA-sequencing data set as a matrix with a row for each unique gene measured and a column for each unique sample in the experiment. This data is typically provided as integer read counts that correspond to the number of times an observed RNA sequence read was aligned to a specific gene transcript. These read count data were pre-processed by first performing quantile normalization to reduce bias between samples, then a log_2_ transformation was applied, which brings large values closer to the distribution and allows for similar variation across different magnitudes. The goal of these pre-processing steps was to reduce systematic bias and improve the analysis of biologically meaningful data [36].

Protein-protein interaction (PPI) graph topology was defined based on protein interaction data from Pathway Commons (https://www.pathwaycommons.org) [13]. These interaction data were further processed by a sparsification technique described in [1], which removed likely false-positive edges. As such, we directly utilized the same interaction network as [1, 2], downloaded from the SCENT Github repository (https://github.com/aet21/SCENT, file: data/net13Jun12.Rda). This PPI network contained 8434 nodes corresponding to unique genes and 303,600 edges describing protein-protein interactions between pairs of nodes.

From here, a (pre-processed) gene expression matrix *D* measuring expression of *M* genes across *N* samples (i.e. *D* is a matrix of dimension *M* × *N*) was overlaid onto the PPI network. First, all nodes corresponding to genes not measured in the gene expression dataset were removed and subsequently only the largest strongly connected component of the resulting network was considered. Next, for each individual sample *k*, taking the corresponding column vector 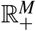 of gene expression, *E* ≔ *D._k_* (i.e., 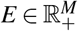), node weight was directly defined as the (pre-processed) gene expression value,

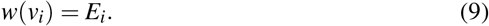

Edge weights were then defined based on the mass-action law [14, 37, 38], which states the rate of interaction is directly proportional to the concentration (or expression) of each gene,

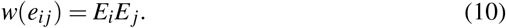

We note that across multiple samples, the same PPI topology was used and new node and edge weights were defined based on the gene expression profile of a given sample.

### 3.2 Gene Network Curvature Analysis

For each individual sample, gene expression was overlaid onto the PPI network and node weights and edge weights were defined as above. A stochastic matrix *P* was created by normalizing edge weights at each node by the sum of outgoing edge weights, such that the sum of outgoing edge weights in *P* sum to 1 for any node,

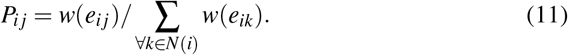

This matrix *P* is notably similar to the transition matrix of a Markov chain, in which the transition probabilities are proportional the the expression level of neighboring nodes. The stationary distribution *π* of the graph, which represents the equilibrium of a random walk on the graph and satisfies *π* = *Pπ* (i.e. *π* is an eigenvector of the stochastic matrix *P* with an eigenvalue of 1) was taken as the normalized first eigenvector of *P*.

Using the normalized edge weights defined by *P*, we applied the combinatorial Forman-Ricci curvature formula, in Eq. (5), to compute curvature at each edge. Then, edge curvature values were used to compute nodal curvature values using Eq. (7). A global average of curvature was computed for each sample, using the stationary distribution *π* to calculate a weighted average over the nodal curvatures, as in Eq. (8). To compare entropy to curvature, the stochastic matrix *P* was used to compute local Shannon entropy at each node and subsequent global entropy rate by Eq. (3).

For local curvature analysis, we develop a differential curvature analysis based on the principles of differential expression analysis [39]. For each individual gene, local curvature values across all samples within a group (i.e. cancer or non-cancer samples) were compared to define a log2 fold-change in curvature (log2FC = log_2_ (mean_2_/mean_1_)) and statistically assessed using an unpaired t-test, with the resulting p-values adjusted for multiple comparisons using the Benjamini-Hochberg step-up procedure [40] to produce false discovery rates (FDR, or q-values). To select genes with significant changes in curvature among the groups, we apply a cutoff of q<0.05 and absolute log2FC>2. These genes with differential curvature were fed into Reactome pathway analysis [41], which utilizes a hypergeometric test to determine pathway overrepresentation, to produce a list of pathways that were retained if the Reactome FDR was q<0.05.

## 4 RESULTS

We now present results on few experiments that illustrate the viability of utilizing gene network Forman-Ricci curvature as an indicator for cellular pluripotency as well as pathway robustness. The aim of this results section is to provide proof-of-concept and is by no means a complete analysis. This said, the results herein lay a foundation for future work, in which we aim to examine higher dimensional structures that more accurately capture biological differentiation dynamics. As such, for the purposes of demonstrating our approach of calculating Forman-Ricci curvature on biological gene networks, we focused our analyses on two previously published single-cell RNA sequencing datasets, first from a stem cell differentiation study [42], and second from a melanoma study including cancer and non-cancer cells [43]. This data is publicly available through the NCBI GEO portal (https://www.ncbi.nlm.nih.gov/geo) under accession numbers GSE75748 and GSE72056, respectively.

### 4.1 Differentiation Cell Type (GSE75748)

We focused on a subset of the stem cell dataset examining a cell-type experiment, which measured gene expression on *n*=1018 single cells from 6 different cell types representing increasingly differentiated cells derived from undifferentiated human embryonic stem cells [42]. Cell-type abbreviations: *hESC* human embryonic stem cell, *NPC* neuronal progenitor cell, *DEC* definitive endoderm cell, *TB* trophoblast-like cell, *HFF* human foreskin fibroblast, *EC* endothelial cell.

We assessed gene network Forman-Ricci curvature and entropy for each individual single-cell sample (Fig. 2). We observed an overall decrease in curvature in more differentiated cells compared to stem and progenitor cells. In agreement with previous studies examining network entropy on this same data [2], a similar trend was observed for entropy. Notably, we observed curvature on these samples to exhibit strongly negative values, which indicated the local edge curvatures on average were predominantly negative, whereas entropy values were normalized to the interval [0,1]. We emphasize the magnitudes of individual curvature values are not nearly as important as the *changes* in curvature between cell types as these changes reflect relative differences in pluripotency. Pearson correlation between curvature and entropy values was 0.9287, suggesting a strong relationship of the two values as expected from the theoretical correlation of entropy and curvature.

**Fig. 2.**
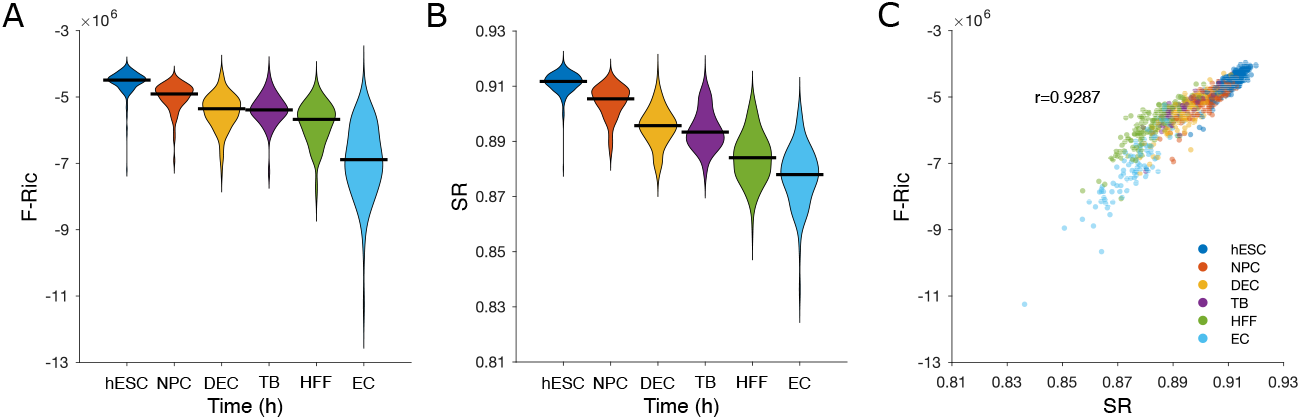
Stem-cell differentiation. A: Violin plot showing distributions of global average curvature in differentiated cell types. Horizontal black bars are medians. A trend of decreasing curvature with increasingly differentiated cell types is observed. B: Same as A but for global entropy. A similar trend of decreasing entropy is observed. C: Scatter plot of global weighted-average nodal curvature vs global entropy. The two quantities were strongly correlated at r=0.9287 (Pearson).

### 4.2 Melanoma (GSE72056)

The melanoma dataset contained *n*=4513 total single-cells from 19 melanoma patients, of which *n_N_*=3256 were non-cancer (normal) cells and *n_T_*=1257 were cancer (tumor) cells [43].

We again computed Forman-Ricci curvature for each single-cell sample (Fig. 3), observing a clear trend of increased curvature in cancer cells compared to non-cancer cells within each patient and across all patients. These findings indicate increased cellular pluripotency or “stemness” in cancer relative to non-cancer tissue.

**Fig. 3.**
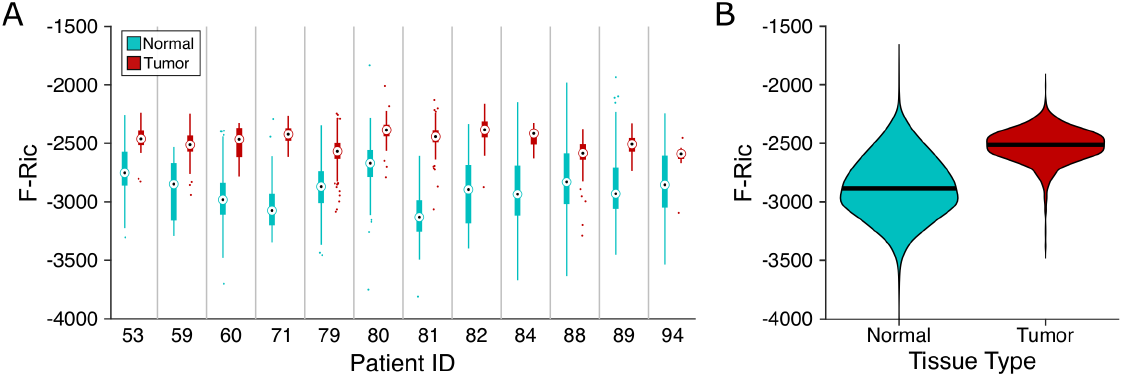
Melanoma. A: Box plot of global average curvature distributions for normal and tumor cells of individual patients. An increase in curvature among tumor cells relative to normal cells was observed in each patient. B: Violin plot of global average curvature in normal and tumor cells across all patients. Horizontal black bars are medians.

We extended this analysis to examine the local curvature at each node (gene), specifically applying a differential curvature analysis approach in order to identify genes with changing curvature between cancer and non-cancer in these melanoma cases. We identified 210 genes with significantly increased curvature and 250 genes with decreased curvature. Subsequently, we utilized Reactome pathway analysis [41] to determine gene pathways to which these increasing and decreasing curvature genes correspond. We found 13 increased curvature pathways including the CCT/TriC protein folding pathway, FGFR4 pathway (oncogene in melanoma [44]), and matrix remodeling pathways, each of which having been implicated in various aspects of melanoma pathogenesis [44–46]. We also found 22 decreased curvature pathways including the FGFR2 pathway (tumor suppressor in melanoma [44]) and several immune pathways, which would typically have protective effects against melanoma [47, 48]. Therefore, the observations of increased curvature in oncogenic pathways and decreased curvature in tumor suppressive pathways highlights the relationship of Ricci curvature and robustness, suggesting that cancer may exhibit increased curvature and thus robustness in pathways beneficial to the cancer. These results demonstrate the capacity of Ricci curvature to identify gene pathway functionality in pathologic processes such as cancer.

## 5 CONCLUSIONS

In this work, we have presented Forman-Ricci curvature as a discrete geometric quantity to evaluate biological gene networks. We were motivated by previous work examining gene network entropy, which found global entropy to be a potential descriptor of differentiation potential [1, 2, 14]. We extended theoretical findings linking entropy and Ricci curvature to establish a relationship between the statistical quantity of entropy and geometric quantity of Ricci curvature [15–17]. In accomplishing this, we selected Forman-Ricci curvature as a discretization of Ricci curvature, owing to its flexibility in assessing more general, high-order biological graph structures. Our findings indicate that Ricci curvature decreases upon cellular differentiation and increases in cancer, similarly to entropy. While network entropy has been previously shown to reflect cellular pluripotency, to our knowledge this is the first example of Ricci curvature representing the same information. Additionally, we demonstrate local analysis of Ricci curvature on a melanoma dataset to reveal several gene pathways with known relevance to melanoma, which exhibits the capacity of Ricci curvature to reveal changes in robustness related to pathway functionality in pathologic processes such as cancer.

While this approach is promising, there are caveats. Many specific details in the gene network analysis are subject to consideration, such as the pre-processing technique, the PPI network topology, and the definitions of node and edge weights. Future directions of this work should include extending the approach to higher-dimensional graph structures, in particular to 2-dimensional simplicial complexes [26], as well as exploring geometric flows such as Ricci flow on biological networks [25]. These avenues could reveal additional biological meaning in regards to cellular pluripotency and differentiation trajectories.

## Acknowledgements

K.M. and R.S. were supported by the National Science Foundation grant ECCS-1749937, the U.S. Air Force Office of Scientific Research grant FA9550-18-1-0130, and E.S. was supported by the German-Israeli Foundation grant I-1514-304.6/2019.

## References

1. C.R. Banerji et al., “Cellular network entropy as the energy potential in Waddington’s differentiation landscape,” Scientific reports, vol. 3, no. 1, pp. 1–17, Oct 2013.

2. A.E. Teschendorff and T. Enver, “Single-cell entropy for accurate estimation of differentiation potency from a cell’s transcriptome,” Nature communications, vol. 8, no. 1, pp. 1–15, June 2017.

3. G. Schiebinger et al., “Optimal-transport analysis of single-cell gene expression identifies developmental trajectories in reprogramming,” Cell, vol. 176, no. 4, pp. 928–943, Feb 2019.

4. C. Weinreb, A. Rodriguez-Fraticelli, F.D. Camargo, A.M. Klein, “Lineage tracing on transcriptional landscapes links state to fate during differentiation,” Science, vol. 367, no. 6479, Feb 2020.

5. B.D. MacArthur, A. Ma’ayan, I.E. Lemischka, “Systems biology of stem cell fate and cellular reprogramming,” Nature reviews Molecular cell biology, vol. 10, no. 10, pp. 672–681, Oct 2009.

6. H. Kitano, “Towards a theory of biological robustness,” Molecular Systems Biology, vol. 3, p. 137, Sep 2007.

7. H. Kitano, “The theory of biological robustness and its implication in cancer,” Systems Biology, pp. 69–88, 2007.

8. I. Amit, R. Wides, Y. Yarden, “Evolvable signaling networks of receptor tyrosine kinases: relevance of robustness to malignancy and to cancer therapy,” Molecular systems biology, vol. 3, no. 1, p. 151, Dec 2007.

9. C. H. Waddington, The Strategy of Genes. London: Allen & Unwin, 1957.

10. J. Wang, K. Zhang, L. Xu, and E. Wang, “Quantifying the Waddington landscape and biological paths for development and differentiation.” Proceedings of the National Academy of Sciences, no. 20, pp. 8257–8262, May 2011.

11. Y.R. Wang, H. Huang, “Review on statistical methods for gene network reconstruction using expression data,” Journal of theoretical biology, vol. 362. pp. 53–61, Dec 2014.

12. T. Klingström and D. Plewczynski D, “Protein-protein interaction and pathway databases, a graphical review,” Briefings in bioinformatics, vol. 12, no. 6, pp.702–713, Nov 2011.

13. I. Rodchenkov et al., “Pathway Commons 2019 Update: integration, analysis and exploration of pathway data,” Nucleic acids research, vol. 48, no. D1, pp. D489–497, Jan 2020.

14. A.E. Teschendorff, P. Sollich, R. Kuehn, “Signalling entropy: A novel network-theoretical framework for systems analysis and interpretation of functional omic data,” Methods, vol. 67, no. 3, pp. 282–293, June 2014.

15. M.K. von Renesse and K.T. Sturm, “Transport inequalities, gradient estimates, entropy and Ricci curvature,” Communications on pure and applied mathematics, vol. 58, no. 7, pp. 923–940, July 2005.

16. K.T. Sturm, “On the geometry of metric measure spaces,” Acta mathematica, vol. 196, no. 1, pp. 65–131, 2006.

17. J. Lott and C. Villani, “Ricci curvature for metric-measure spaces via optimal transport,” Annals of Mathematics, vol. 169, no. 3, pp.903–991, May 2009.

18. M. Pouryahya, J.H. Oh, J.C. Mathews, J.O. Deasy, A.R. Tannenbaum, “Characterizing cancer drug response and biological correlates: a geometric network approach,” Scientific reports, vol. 8, no. 1, pp.1–2, Apr 2018.

19. M.P. do Carmo, Riemannian Geometry. Boston, MA: Birkhäuser, 1992.

20. Y. Ollivier, “Ricci curvature of Markov chains on metric spaces,” Journal of Functional Analysis, vol. 256, no. 3, pp. 810–864, Feb 2009.

21. W.V.D. Hodge, The Theory and Applications of Harmonic Integrals, 2nd ed. Cambridge, UK: Cambridge University Press, 1952.

22. R. Forman, “Bochner’s method for cell complexes and combinatorial Ricci curvature,” Discrete and Computational Geometry, vol. 29, no. 3, pp. 323–374, Feb 2003.

23. R.P. Sreejith, K. Mohanraj, J. Jost, E. Saucan, A. Samal, “Forman curvature for complex networks,” Journal of Statistical Mechanics: Theory and Experiment, vol. 2016, no. 6, p. 063206, June 2016.

24. E. Saucan, R.P. Sreejith, R.P. Vivek-Ananth, J. Jost, A. Samal, “Discrete Ricci curvatures for directed networks,” Chaos, Solitons & Fractals, vol. 118, pp. 347–360, Jan 2019.

25. M. Weber, E. Saucan, J. Jost, “Characterizing complex networks with Forman-Ricci curvature and associated geometric flows,” Journal of Complex Networks, vol. 5, no. 4, pp. 527–550, Aug 2017.

26. E. Saucan, M. Weber, “Forman’s Ricci curvature-from networks to hypernetworks,” In Proc. International conference on complex networks and their applications, Dec 2018, pp. 706–717.

27. R. Sandhu et al., “Graph curvature for differentiating cancer networks,” Scientific reports, vol. 5, no. 1, pp. 1–13, July 2015.

28. R. Sandhu, S. Tannenbaum, T. Georgiou, A. Tannenbaum, “Geometry of correlation networks for studying the biology of cancer,” In Proc. IEEE 55th Conference on Decision and Control (CDC) 2016, Dec 2016, pp. 2501–2506.

29. M. Weber, J. Stelzer, E. Saucan, A. Naitsat, G. Lohmann, J. Jost, “Curvature-based methods for brain network analysis,” arXiv preprint, arXiv:1707.00180, July 2017.

30. H. Farooq, Y. Chen, T.T. Georgiou, A. Tannenbaum, C. Lenglet, “Network curvature as a hallmark of brain structural connectivity,” Nature communications. vol. 10, no. 1, pp. 1–11, Oct 2019.

31. C. Wang, E. Jonckheere, R. Banirazi, “Wireless network capacity versus Ollivier-Ricci curvature under Heat Diffusion (HD) protocol,” In Proc. IEEE 2014 American Control Conference, 2014, pp. 3536–3541.

32. R. Sandhu, T. Georgiou, A. Tannenbaum, “Market fragility, systemic risk, and Ricci curvature,” arXiv preprint, arXiv:1505.05182, May 2015.

33. R. Sandhu and J. Liu, “Maxwell’s Demon: Controlling entropy via discrete Ricci flow over networks,” In Proc. International Conference on Network Science, Springer, Cham, Jan 2020, pp. 127–138.

34. L.A. Demetrius, “Boltzmann, Darwin and directionality theory,” Physics reports, vol. 530, no. 1, pp. 1–85, Sep 2013.

35. M.L. Siegal, D.E. Promislow, A. Bergman, “Functional and evolutionary inference in gene networks: does topology matter?” Genetica, vol. 129, no. 1, pp. 83–103, Jan 2007.

36. X. Liu et al., “Normalization methods for the analysis of unbalanced transcriptome data: a review,” Frontiers in bioengineering and biotechnology, vol. 7, p. 358, Nov 2019.

37. C.M. Guldberg and P. Waage, “Uber die chemische Affinität,” J. prakt. Chem, vol. 19, no. 69, p. 13, 1879.

38. G. Schreiber, “Protein-protein interaction interfaces and their functional implications,” in Protein-protein interaction regulators, Royal Society of Chemistry, pp.1–24, Dec 2020.

39. J. Costa-Silva, D. Domingues, F.M. Lopes, “RNA-Seq differential expression analysis: An extended review and a software tool,” PloS one, vol. 12, no. 12, p. e0190152, Dec 2017.

40. Y. Benjamini, Y. Hochberg, “Controlling the false discovery rate: A practical and powerful approach to multiple testing,” Journal of the Royal Statistical Society: Series B, vol. 57, no. 1, pp. 289–300, Jan 1995.

41. B. Jassal et al., “The reactome pathway knowledgebase,” Nucleic Acids Res., vol. 48, no. D1, pp. D498–D503, Jan 2020.

42. L.F. Chu et al., “Single-cell RNA-seq reveals nodel regulators of human embryonic stem cell differentiation to definitive endoderm,” Genome biology, vol. 17, no. 1, pp. 1–20, Dec 2016.

43. I. Tirosh et al., “Dissecting the multicellular ecosystem of metastatic melanoma by singlecell RNA-seq,” Science, vol. 352, no. 6282, pp.189–196, Apr 2016.

44. M. Czyz, “Fibroblast growth factor receptor signaling in skin cancers,” Cells, vol. 8, no. 6, p. 540, June 2019.

45. C. Boudiaf-Benmammar, T. Cresteil, R. Melki, “The cytosolic chaperonin CCT/TRiC and cancer cell proliferation,” PloS one, vol. 8, no. 4, p. e60895, Apr 2013.

46. S. Napoli et al., “Functional roles of matrix metalloproteinases and their inhibitors in melanoma,” Cells, vol. 9, no. 5, p. 1151, May 2020.

47. A. Passarelli, F. Mannavola, L.S. Stucci, M. Tucci, F. Silvestris, “Immune system and melanoma biology: a balance between immunosurveillance and immune escape,” Oncotarget, vol. 8, no. 62, p. 106–132, Dec 2017.

48. M. Tucci et al., “Immune system evasion as hallmark of melanoma progression: the role of dendritic cells,” Frontiers in oncology, vol. 9, p. 1148, Nov 2019.

